# Elucidating TolC Protein Dynamics: Structural Shifts Facilitate Efflux Mediated β-lactam Resistance

**DOI:** 10.1101/2024.02.22.581573

**Authors:** Isik Kantarcioglu, Ilona K. Gaszek, Tandac F. Guclu, M. Sadik Yildiz, Ali Rana Atilgan, Erdal Toprak, Canan Atilgan

## Abstract

Efflux-mediated β-lactam resistance represents a significant public health challenge, limiting the efficacy of various β-lactam antibiotics against numerous clinically relevant pathogenic bacteria. Structural and functional analyses have revealed that the efflux protein TolC in several Gram-negative bacteria serves as a conduit for antibiotics, bacteriocins, and phages, affecting bacterial susceptibility and virulence. In this study, we conducted a comprehensive examination of the efflux of β-lactam drugs mediated by TolC, employing extensive experimental and computational analyses. Our computational investigations into the molecular dynamics of drug-free TolC revealed critical unidirectional movements of the trimeric TolC and identified residues significantly involved in TolC opening. To corroborate these findings, we performed a whole-gene-saturation mutagenesis assay, systematically mutating each residue of TolC to 19 other amino acids and measuring the fitness effects of these mutations under β-lactam-induced selection. The β-lactams oxacillin, piperacillin, and carbenicillin were selected for this study because they are effluxed by the AcrAB-TolC complex with varying efficiencies. This approach clarified the similarities and differences in the efflux processes of the three β-lactam antibiotics through the trimeric TolC. Further analysis of TolC’s efflux mechanism for these β-lactam antibiotics via steered molecular dynamics simulations revealed the existence of general and drug-specific mechanisms employed by TolC. We identified key positions at the periplasmic entry of TolC whose altered dynamics influence long-range efflux motions as allosteric modulators. Our findings provide valuable insights into the structural dynamics of TolC, establishing a foundation for understanding the key mechanisms behind multidrug resistance and principles for designing new antibiotics and antibiotic derivatives capable of circumventing the bacterial efflux mechanism.

## INTRODUCTION

### TolC is an outer membrane protein with a critical role in antibiotic efflux

Gram-negative bacteria have evolved transport mechanisms to move macromolecules and toxic compounds across membranes (Paulsen et al. 2006). These mechanisms play important roles in drug resistance to current and future antibiotics (Nikaido 1996). Bacterial efflux is one of the primary systems responsible for the emergence of multidrug resistance since efflux machineries in bacteria can pump out several different types of antibiotics with varying rates and specificity (Levy and Marshall 2004; Nikaido 2009). The Resistance-nodulation-division (RND) family is one of the major efflux pump superfamilies in Gram-negative bacteria that can efflux a broad range of substrates, including clinically important antibiotics such as β-lactams, bile salts, and detergents (Poole 2002; Nikaido 2011). The major multidrug-resistant efflux pump in *Escherichia coli* (*E. coli*) is the AcrAB-TolC protein complex (Fralick 1996). In *E. coli*, the outer membrane protein TolC can transiently associate with the protein pairs AcrAB, MacAB, and EmrAB to create various tripartite efflux pump assemblies as a response to elevated antibiotic concentrations inside bacteria (Piddock 2006). These efflux assemblies, which span the length of the periplasmic space, expel toxic substances from the cytoplasm to the extracellular region.

TolC is a multifunctional protein that is subject to conflicting evolutionary selection pressures. While TolC is crucial to protect bacterial cells against antibiotics and other antibacterial compounds, its presence can impose a risk for bacterial cells, since TolC can also function as a portal for the transport of bacteriocins and bacteriophages into the cells (**Figure 1A**) (Ayhan et al. 2016). Thus, bacteria need to strike an evolutionary balance between making TolC available when efflux is necessary and limiting it as an entry point when efflux presents risk (Puchta et al. 2016).

**Figure 1.**
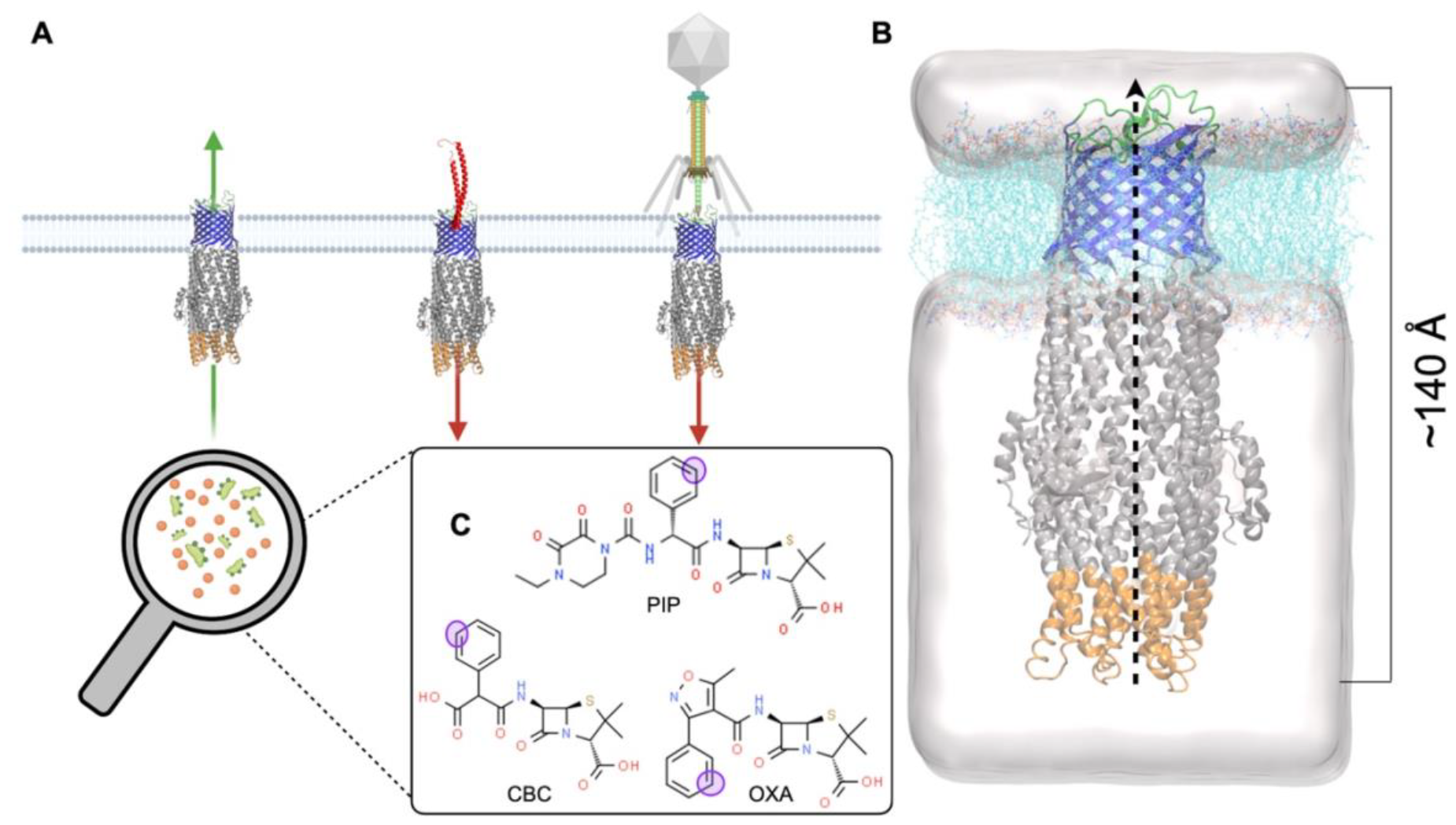
TolC and ligand structures. **(A)** TolC structure positioned in outer membrane. The major known role of TolC in *E. coli* is to expel antibiotics and bile salts to the outside of the cell which is essential for bacterial survival (green arrow). However, it is also an entry point for colicin-E1 (partial structure shown in red) and some bacteriophages which results in cell death (red arrow). **(B)** TolC embedded in a POPE lipid membrane (cyan) and placed in a simulation box shown as a white blob; extracellular loops are colored in green, membrane embedded β-barrel is colored in blue, α-helices extended through periplasmic space are colored in gray and the approximately 30 Å part located at the periplasmic tips of the protein is colored in orange. This coloring scheme is maintained throughout the manuscript. The reaction coordinate of the drug is defined from the entry point to the exit and spans ∼140 Å (dashed line). **(C)** Three-dimensional representations of the antibiotics studied in this work. Carbenicillin (CBC), piperacillin (PIP), and oxacillin (OXA); SMD pulling atoms for the drug molecules are shown as purple circles.

Structural studies reveal that TolC functions as a homotrimer. Its 12-stranded β-barrel part is embedded into the outer membrane and is approximately 40 Å long; its cone-like α−helical coiled-coil domain extends into the periplasm (**Figure 1B**) spanning an additional 100 Å (Koronakis et al. 2000). The outer membrane channel opening, especially periplasmic end opening, is necessary to initiate the process of the expulsion of the antibiotic molecules (Bavro et al. 2008).

### Previous biophysical studies utilizing computational tools uncovered important structural regions and rearrangements in TolC and its homologs

Outer membrane protein OprM from *Pseudomonas aeruginosa* is one of the closest homologs of *E. coli* TolC. Normal mode analysis (NMA) based on elastic network models (Atilgan et al. 2001) was previously used to uncover the conformational states of OprM and how they compare to the open state of TolC (Phan et al. 2010). In the closed form of TolC, coiled-coil helices linked by salt bridges and hydrogen bonds at the tip of the periplasmic entrance provide selectivity and constrictions for substance transport, particularly maintained by D371 and D374 (Bavro et al. 2008; Pei et al. 2011). CmeC found in *Campylobacter jejuni* is another homologue that shares structural similarities with TolC. Substrate efflux mechanisms with CmeC have also been simulated by others (Newman and Khalid 2023) using steered molecular dynamics (SMD) simulations (Izrailev et al. 1997). Their results indicated the anionic bile acid moves up the channel by climbing a ladder of acidic residues that align in the interior surface of the protein. Unlike CmeC, TolC has far fewer charged residues along the efflux path, which indicates a different action mechanism.

These findings and the critical importance of TolC in antibiotic resistance motivate our exploration of the structural shifts in the TolC protein at the atomistic level. We first utilized molecular dynamics (MD) simulations for both the closed and the open forms of drug-free TolC to determine its intrinsic dynamics and structural transitions that are difficult to observe by most experimental methods. Principal component analysis (Jolliffe 2002; Lange et al. 2008) of MD trajectories suggested that TolC has an intrinsic expelling movement in both closed and open forms. The dominant modes of these forms displayed net movement in the direction of efflux and, hence, suggest these modes may play an important role in expelling antibiotic molecules. Since TolC conformations can switch between closed and open forms, we hypothesized that there must be important residues whose perturbation might tip the equilibrium between the two conformations. To systematically search for these residues, we utilized the perturbation response scanning (PRS) that we previously developed as a robust and now a widely used method for identifying essential residues that trigger conformational changes of proteins (Atilgan and Atilgan 2009; Atilgan et al. 2010). PRS relies on linear response theory to determine specific residues that, when perturbed via external forces, might lead to conformational changes in proteins; PRS has been successfully used by several groups to identify structurally important regions not only in many monomeric proteins but also in multimeric assemblies such as HSP90 (Penkler et al. 2018). PRS also was applied to unravel motions that trigger uncoating when applied to a multiprotein capsid comprised of 60 protomeric units, each of which contained four heterogenic subunits (Ross et al. 2018).

### We explored dynamical shifts within the TolC structure that enable the efflux of three β-lactam antibiotics: carbenicillin, piperacillin, and oxacillin, as illustrated in Figure 1C

The choice of β-lactam antibiotics was due to their great importance in fighting against bacterial infections and the rapid evolution of β-lactam resistance in clinical and community settings. Almost 80 percent of all antibiotics used in clinics are β-lactams. Although resistance to β-lactams is often attributed to spread of β-lactamase genes producing β-lactamases that render β-lactams ineffective by hydrolyzing them, many β-lactam antibiotics are ineffective against Gram-negative bacteria because of their fast efflux from the cytoplasm. For instance, we previously showed that disruption of the AcrAB-TolC complex in *E. coli* can reduce the required dose of oxacillin to kill bacterial cells by almost three orders of magnitude (Ayhan et al. 2016). We chose oxacillin, piperacillin, and carbenicillin for our study due to the distinct efflux efficiencies of the AcrAB-TolC complex towards these antibiotics—high for oxacillin, moderate for piperacillin, and low for carbenicillin. To substantiate these efficiencies, we compared the minimum inhibitory concentrations (MICs) needed to inhibit a wild-type *E. coli* strain, a *tolC*-deficient mutant, and the mutant harboring a *tolC*-carrying rescue plasmid, as shown in **Figure 2**. We also systematically explored the fitness effects of single amino acid replacements at each position in TolC using a saturation mutagenesis library under selection with each of these drugs. Finally, we utilized SMD simulations to calculate work curves during passage of these drug molecules through TolC homotrimer opening to determine drug specific and generalist TolC residues involved in β-lactam efflux.

**Figure 2.**
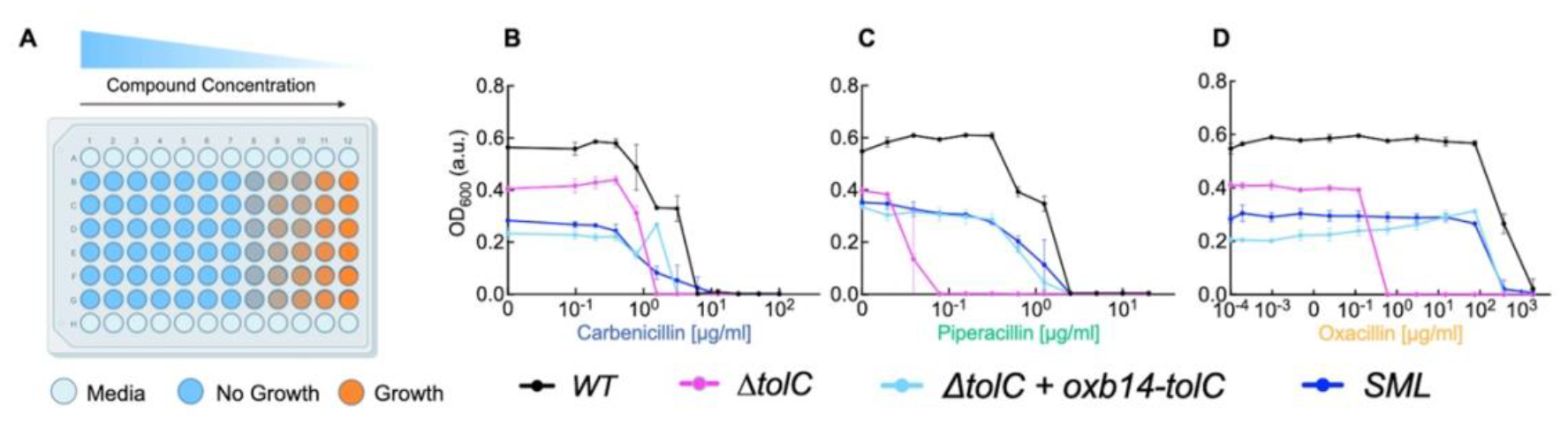
Minimum inhibitory concentration (MIC) measurements. **A**. MIC assay plate setup. Blue wells indicate no growth; color gradation towards orange indicates the level of growth. **(B-D)** Three different antibiotics were used as selection factors. We incubated three strains and the Single Mutation Library (SML, Methods) in the presence of increasing drug concentrations and monitored bacterial growth for ∼18h. We used blue x-axis label for carbenicillin, green x-axis label for piperacillin, and orange x-axis label for oxacillin. This coloring scheme representing the antibiotics was maintained throughout the manuscript. Black line represents dose response curves for wild type (BW25113) *E. coli* strain. Magenta lines represent *E. coli* strain with *tolC* gene deletion (*ΔtolC*). Cyan lines represent *ΔtolC+*pOxb14*tolC*. Blue lines represent SML strain which contains all possible amino acid mutations. Background-corrected OD600 values after 18 hours of incubation measure cell growth on the y-axis, with error bars showing standard deviations across three replicates per concentration. Raw data with background corrections is provided in **Table S1**.

## RESULTS and DISCUSSION

### TolC dynamics are inherently optimized for efflux

We performed MD simulations to gain insights about TolC structural dynamics for closed and open states of the wild type TolC (**Figure 3A**). Root-mean square fluctuations of the residues (RMSFs) were highest at the periplasmic side for all simulated structures (**Figure 3B**). However, the fluctuations were greater on the periplasmic side in the open form, and on the outer membrane side in the closed form, which might be indicative of the changes in the closing/opening motion of the protein (**Figure 3B**). Finally, the larger fluctuations near residue 200 correspond to the equatorial domain helices in the widest region of the protein contacting water and are not functionally relevant. Consequently, their mobility remains the same irrespective of the open/closed forms.

**Figure 3.**
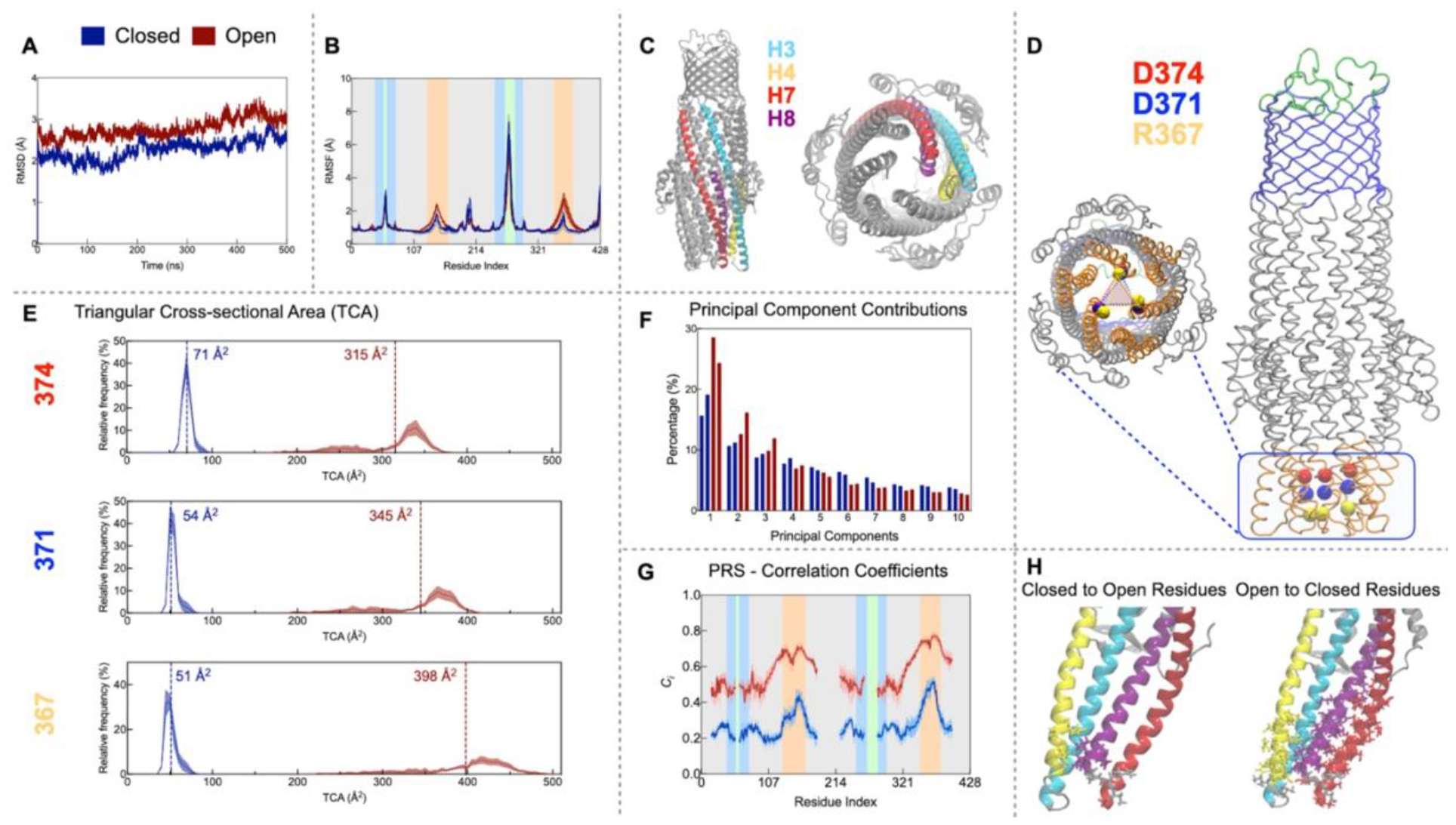
Molecular dynamics of ligand-free TolC. **A**. C_α_ RMSDs of TolC remain in the 2-4 Å range during the 500 ns MD trajectories of both closed (blue), and open (red) forms. **B**. C_α_ RMSFs; colored vertical stripes follow the coloring of the TolC regions **Figure 1B**: green extracellular loops, orange periplasmic entry region, blue β-barrel, gray the rest. Standard error of the mean (SEM) obtained from the average of the three chains in the trimer are shown as shades. **C**. Specific four helices of one TolC protomer highlighted; side and bottom views. **D**. Selected pore constricting trimeric C_α_ atoms visualized as spheres on the protein (R367 yellow, D371 blue, D374 red); the bottom view highlights the triangular shape. **E**. Triangular cross-sectional area (TCA) histograms of residues in D. **F**. Contribution of the top 10 PCs, obtained from two separate trajectory pieces (240-280 ns and 460-500 ns) in each of the closed (blue) and open (red) form simulations. **G**. Correlation coefficients obtained from PRS for closed and open forms. SEM shown as shades. Highest mobility regions excluded in the construction of the correlation matrix to capture motions of the main structure appear as disconnected regions in the graphs. **H**. PRS determines single residues implicated in channel opening mainly located on four helices shown in C. Top-ranked residues for opening (*C*_*i*_>0.50): 160, 366-369, 369-374; top-ranked residues for closing (*C*_*i*_>0.70): 129,132,133,135-137,140,149,150,152-164,341-377. D371 has the highest correlation value of 0.54 and 0.82, respectively for closed and open forms.

The greatest conformational changes were observed in the H7/H8 helices. H3/H4 helices were comparatively stable (**Figure 3C**) (Bavro et al. 2008; Weng and Wang 2019). D371 and D374 on H8 are at the points of the two of the narrowest constrictions (Weng and Wang 2019). In fact, D371 was shown to be important in gating, with increased susceptibility to various substances in the D371V mutant (Vaccaro et al. 2008). R367 is another site that makes strong interactions to hold inter-protomer and intra-protomer helices together (R367-D153 and R367-T152 for the former and Y362-D153 for the latter) (Bavro et al. 2008). Since TolC is a homotrimer, we calculated the triangular cross-sectional area (TCA) using the C_α_ atoms of these specific residues, R367, D371, and D374, which form a ladderlike entry region into the pore used in efflux (**Figure 3D**). TCAs for the open structure are expectedly much larger than the closed form. However, the largest area is at the level of the innermost D374 residue in the closed form while the reverse is true for the open form, i.e., the truncated cone formed by these three residues is reversed during opening (**Figure 3E**).

To understand the underlying motions that collectively orchestrate TolC dynamics in the absence of drugs, we monitored the principal components (PCs) obtained from the trajectories. The top three PCs have similar amounts of contributions to the dynamics in the closed form, while the most collective mode of motion is more separated than the rest in the open form (**Figure 3F**). In its closed form, the primary motion (PC1) induces a torque along the efflux axis in the lower half and a stretching opening in the upper half; the reverse is true for PC2. PC3 is an upward pumping motion (see **Movies S1-S3**). In the open form, motions in the open periplasmic side overwhelm the dynamics, while PC2 and PC3 overlap with the top two PCs of the closed form (see **Movies S4-S6**). Thus, TolC remains primed for efflux even in the absence of drugs, more so when closed than open.

PRS calculations determined the key residues involved in triggering the open-to-closed and closed-to-open conformational changes (**Figure 3G-H**). Our findings show that the closed-to-open correlation coefficients, Ci, are significantly lower than open-to-closed ones, suggesting the protein naturally tends to close from an open state and requires specific residue perturbations for opening from a closed state. Despite the less stringent selection of the threshold we employed, we found far fewer residues that can enforce channel opening. Moreover, residues located on the H7/H8 helices have higher *C*_*i*_ values than the H3/H4 helices, reinforcing the active role of the former pair of helices in channel opening as previously reported (Bavro et al. 2008; Weng and Wang 2019). Aligning with previous determinations, our results confirmed the closed form as TolC’s stable state, upon which we based our continued simulations.

### TolC facilitates drug efflux through dynamic changes in a critical interaction network

We used steered molecular dynamics (SMD) simulations to track the work produced as antibiotics move through TolC (**Figure 4A**). We calculated the average work to transport from the periplasmic to the extracellular side, with the order being carbenicillin > piperacillin > oxacillin. We also monitored the drug-protein interactions along this path (**Figure 4B-D**). We found common interactions for all antibiotic molecules we tested at the extracellular side during the final exit of the antibiotic (those in the green stripe region in **Figure 4**). However, carbenicillin formed more bonds than the other antibiotics, correlating with our findings that its efflux rate through E. coli TolC is lower. The rise in hydrogen bonds during carbenicillin’s transfer increased energy expenditure, prompting us to explore if this was due to direct contacts with TolC. Our analysis revealed that the average number of hydrogen bonds with TolC’s pore lining and the derivative of average work shared similar patterns, albeit with a slight shift on the reaction coordinate (**Figure 4B**). This shift results from basing the reaction coordinate on the pulling atom (purple in **Figure 1C**), in contrast to counting hydrogen bonds from any part of the drug molecule and TolC.

**Figure 4.**
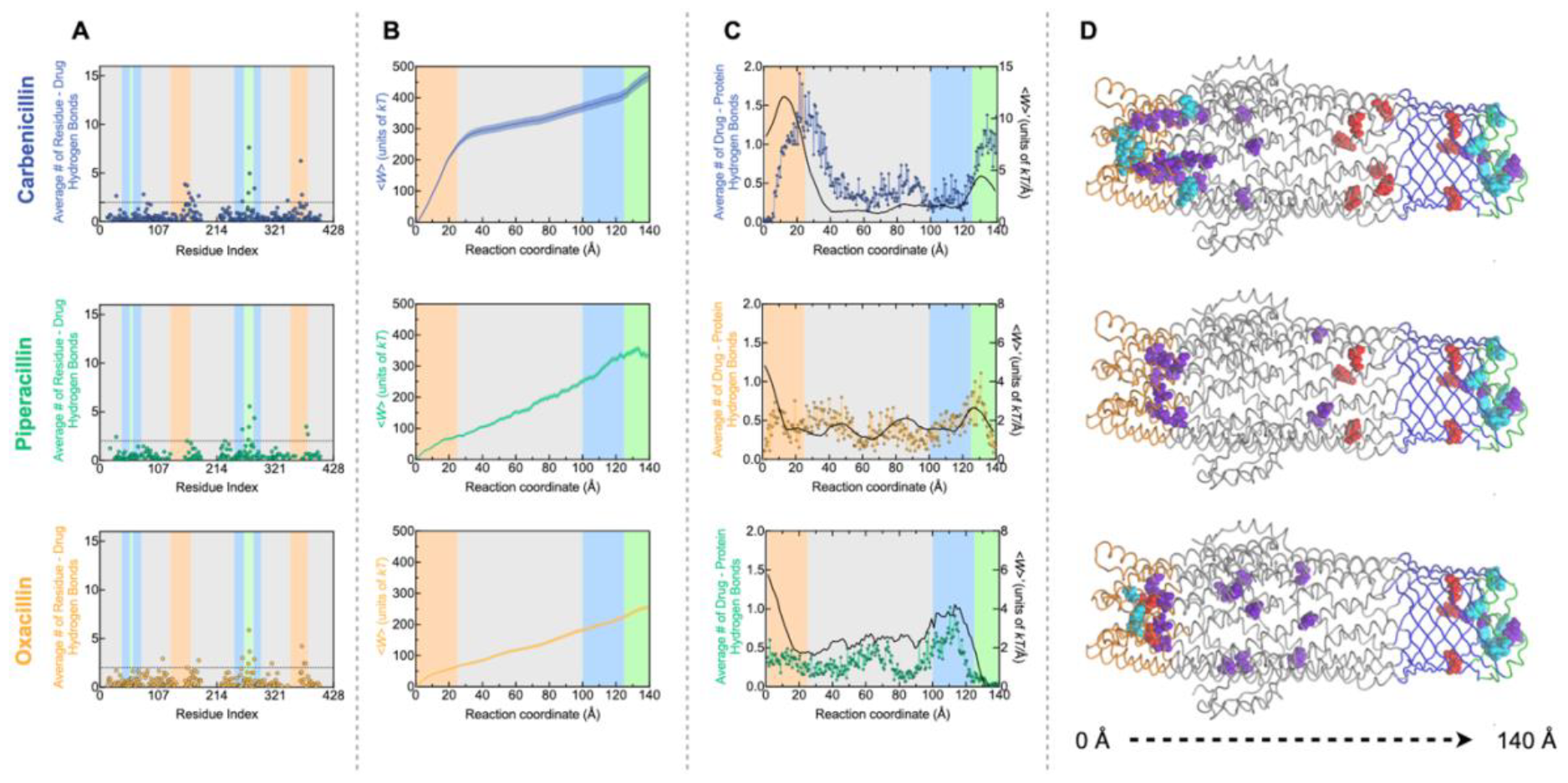
Mapping drug-TolC interactions. Carbenicillin (blue; top row), piperacillin (green; middle row), oxacillin (yellow; bottom row). Vertical stripes use the color code of the regions of TolC depicted in **Figure 1B. A**. The work curves from SMD simulations. **B**. The average number of hydrogen bonds between drug and TolC along the reaction coordinate (colored; left y-axis) and the first derivative of the work (black; right y-axis). **C**. Residues that have more than an average of two hydrogen bonds (above dashed line in **D**) plotted on the 3D representation of the protein; charged residues red, polars purple, and hydrophobics cyan. **D**. Number of hydrogen bonds formed between a given residue and the drug averaged over 45 SMD simulations.

Residues that formed two or more hydrogen bonds with selected antibiotics were mainly clustered at the periplasmic tip of the protein (**Figure 4C**). For carbenicillin, the polar residues lining the pore played an important role in retarding the progress of the drug movement. Moreover, the hydrophobics at the entry region seemed to form a barrier for carbenicillin and piperacillin but not for oxacillin. The lack of this initial barrier might be the main reason behind the efficient efflux of oxacillin and why it is a nonviable drug for treating Gram-negative infections (**Figure 2 and Figure S1**). Once the antibiotic molecules overcame the resistance at the TolC entrance region marked by the orange regions in **Figure 4**, they accessed the membrane bound part of the pore with relative ease.

We next looked more carefully at the movement of carbenicillin through the protein channel since it experiences a larger number of interactions at the entry to the protein than the other antibiotics and forms extra hydrogen bonds with charged residues in the extracellular barrel region (suggested by the middle peak in **Figure 4B**). Supplementary Video S7 illustrates a standard path of carbenicillin. While carbenicillin was moving through the first ∼30 Å long region, the triplet salt bridges between R367 and D153 were typically broken. Interestingly, a permanent salt interaction with D371 is established instead (**Figure S2**). Here, the negatively charged carbenicillin competed with D153/D371 on the H2/H7 helices. Once carbenicillin moved beyond this region, the drug molecule moved predominantly unhindered through the next 90 Å. In fact, the charged and polar residues aided this last part of the journey by making a hand-over-hand type of coordinated motion that advanced the drug to the outer region of the channel. Finally, in the last ∼25 Å, the drug overcame the hydrophobic network of interactions that seal the exit before it finally left the channel. The latter motion is common to all drugs, as exemplified in **Videos S8 and S9**.

### Deep mutational scanning of TolC indicates mutations that confer antibiotic sensitivity disrupt the dynamical interaction network and hinder efflux

Using a sequencing-based selection assay, we quantified the fitness impact of single amino acid substitutions across TolC, with carbenicillin, piperacillin, and oxacillin as selection agents. Most positions in TolC showed optimized efflux activity, with few exhibiting notable fitness deviations (**Figure 5A; Figures S3-S5**). Fifty-one amino acid level replacements in the 428 residues were recorded. Eight mutations were common under all selection conditions; all were sensitivity conferring (**Figure 5A-B**). Consistent with the observation that the structure of TolC is less optimized for carbenicillin efflux than for efflux of the other antibiotics, increasing carbenicillin selection strength from 10 μg/ml to 20 μg/ml resulted in a decrease in the number of resistance-conferring mutations and a simultaneous increase in the number of sensitivity-conferring mutations at the periplasmic region of the protein. Nevertheless, the fitness effects of mutations under different carbenicillin concentrations for selection were highly correlated; of the 27 and 32 mutations recorded in the respective concentrations, 23 were common (**Figure 5A, Figure S5**).

**Figure 5.**
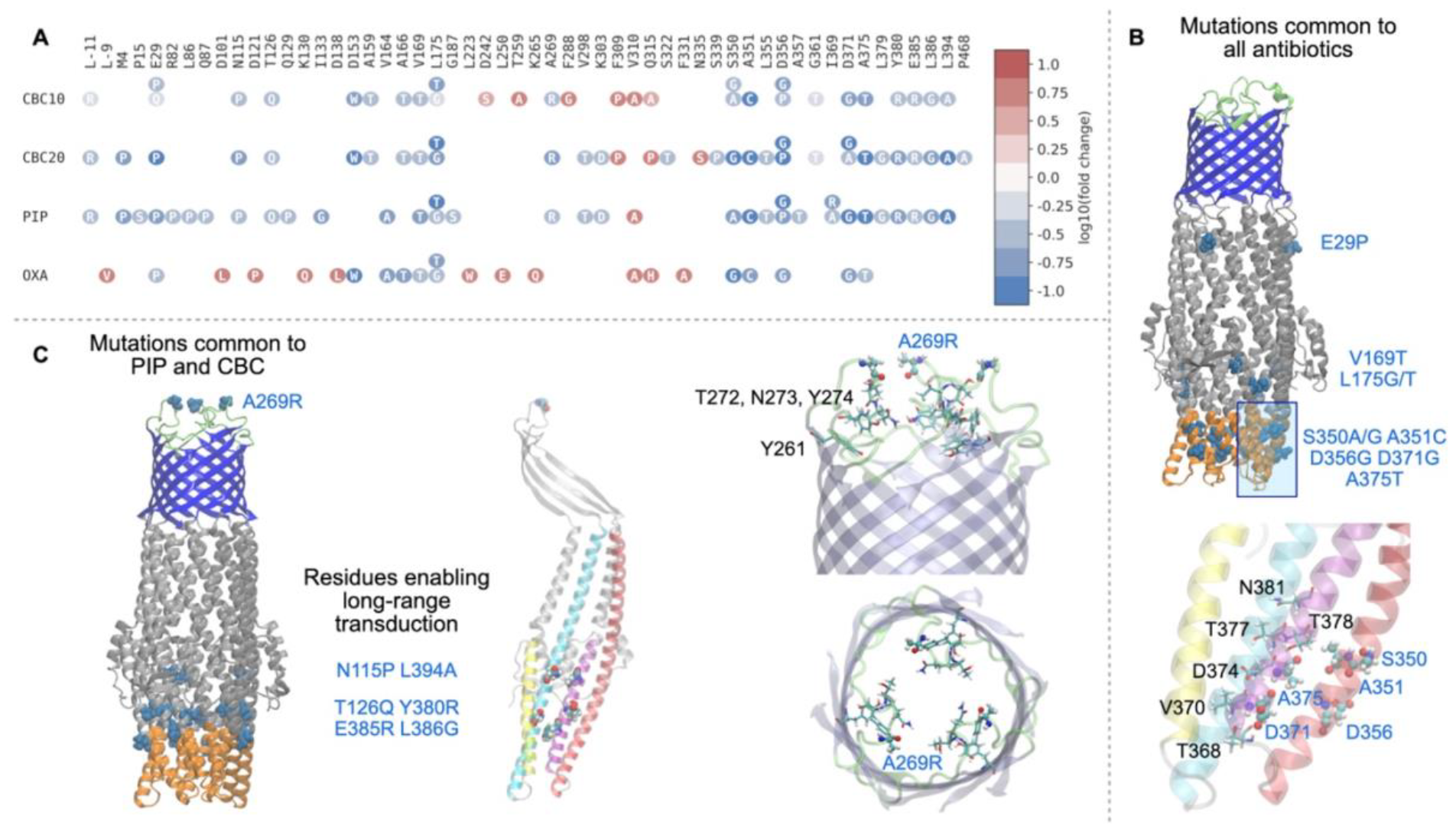
Single mutant library selections reveal the TolC residues critical for efflux activity. **A**. Abacus plots summarizing TolC replacements that lead to significant changes in fitness values under selection of carbenicillin (10 μg/ml; CBC10), carbenicillin (20 μg/ml; CBC20), piperacillin (20 μg/ml; PIP), and oxacillin (460 μg/ml; OXA). Excluding synonymous mutations affecting sensitivity, fitness values are represented by a color spectrum from dark blue (more susceptible) to dark red (more resistant). **B**. Mutated residues identified in all library selections, which confer sensitivity, are highlighted on the protein structure. Below, mutations at the periplasmic entry region are depicted on a single protomer (coloring of **Figure 3C** for the helices is used), with sensitivity-conferring amino acids in CPK representation and labeled in blue. Residues with a high number of antibiotic interactions in SMD simulations (**Figure 4D**) are shown in licorice representation and labeled in black. **C**. Mutations shared by carbenicillin, and piperacillin are illustrated on the protein structure. In the center, residues near the equatorial region enabling long-range transduction along the major helices are shown on a single protomer with the coloring in **Figure 3C**. On the right, extracellular exit site residues are depicted in both side and top views. Residues with high antibiotic interaction counts from SMD simulations are displayed in licorice representation and labeled in black, while the hydrophobic extracellular loop region is shown in green with A269 highlighted in CPK.

The mutations common to all selection conditions were clustered in two regions of the protein: S350A/G, A351C, D356G, D371G, and A375T are all near the periplasmic entry region, while the hydrophobic to polar changes V169T and L175G/T occur near the equatorial domain slightly up the channel (the remaining E29P mutation might destabilize protein folding). Sensitivity-conferring mutations clustered at the periplasmic entrance can disrupt the interactions between the adaptor protein and TolC, and prevent the operation of the efflux machinery. Another potential function of these mutations is to enhance the interactions with applied antibiotics and, hence, impede the efflux of antibiotics into the extracellular space (**Figure 5B, bottom**). When we looked at the specifics of how the mutations depended on the selection agents, we found that evolution again exploits the network of local interactions utilized in efflux. Oxacillin, which has low efficacy against *E. coli* because it is highly effluxable by the AcrAB-TolC, leads to the selection of the least number of deviations and potentially the development of further resistance (**Figure 5A**). Conversely, the efflux pump had fewer positions that are prone to developing resistance to carbenicillin and piperacillin; most mutations appearing in these backgrounds led to elevated susceptibility to these drugs. In fact, these drugs have seven additional common mutations (**Figure 5C**): those mutations near the equatorial region (N115P and L394A) and those in the region right below (T126Q, Y380R, E385R, L386G) are all in positions where the TolC-carbenicillin/piperacillin interactions are lowest (**Figure 5A and C**) and are in the rigid domains (low RMSFs; **Figure 3B**). We conjecture that these residues occupy important positions that act as the transducers of the intrinsic efflux motions of TolC reflected in the most dominant PCAs (**Movie S1-S3**). Any mutation that ‘softens’ this rigidity is expected to alter the efflux mechanism and lead to sensitivity.

At the final exit region, extensive interactions between the drugs and polar/charges residues Y261, T272, N273, Y274 are critical to present the drug to the opening of the portal made up of the flexible hydrophobic patch of residues forming the loop, i.e., G268, A269, A270, G271 (**Figure 5C, right**). The hydrophobic to polar mutation A269R, which confers sensitivity to both carbenicillin and piperacillin, disrupts the hydrophobic balance near the exit region. This may result from redistributing the network of drug-TolC interactions that contribute to the increased amount of work required to expel the charged drug out of this region. We note that this change would be an entropy dominated shift that involves number and distribution of water molecules near the exit site.

### The D371G mutation reveals an allosteric control mechanism spanning the entire efflux pump

Since TolC is optimized for efflux, we tested whether and how these mutations might lead to allosteric effects along with the disrupted local interactions. We selected the D371G mutation that commonly emerged as a sensitivity-conferring mutation in all four of our saturation mutagenesis library selection assays (**Figure 5A**). This is counterintuitive since the replacement of aspartate with glycine, a smaller amino acid with no charge, might be expected to increase the volume available to these negatively charged drugs and make it easier for them to be carried away from the initial entry region. D371 is also the top residue providing the highest correlation coefficient in manipulating both the closed-to-open and open-to-closed conformational change in our PRS analysis (**Figure 3H**).

We conducted 45 SMD simulations on the D371G mutant with carbenicillin and found the energy required for efflux did not change compared to the wildtype TolC. However, we noted both a minor translation in the curve in the entry region (**Figure 6A-C**) and a substantial reduction in hydrogen bonding partners there (**Figure 6C**). Drug-free MD simulations showed that the substitution D371G eliminates likely dynamic salt bridges with R367 on the neighboring trimeric chain (Koronakis et al. 2000), (Wang et al. 2017).

**Figure 6.**
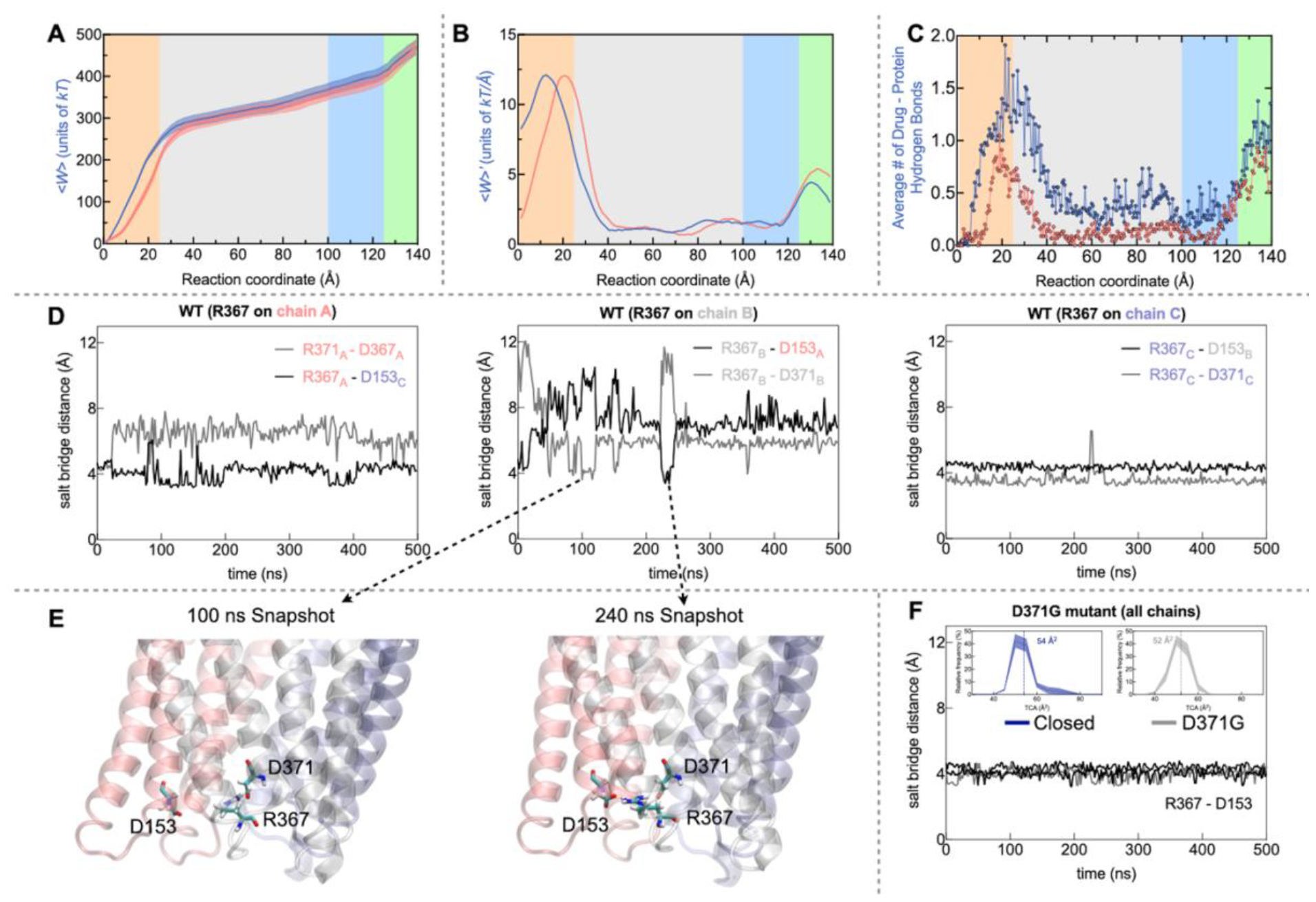
The global and local effects of the D371G mutation on TolC. **A-C**. Carbenicillin SMD findings comparing wild type and D371G mutant. **D**. Dynamical shifts in R367 salt bridges on the three chains of wild type TolC in the absence of drugs. Manipulation of R367-D153 salt bridge is necessary to provide flexibility for drug entry and initial efflux (see also **Figure S2**). R367 has a canonical salt bridge with the D153 on the adjacent chain in the heterotrimer. However, D371 residing on the same chain tends to compete with D153 for this salt bridge. The time evolution of the relevant distances is shown for each chain. **E**. Possible R367 side chain positions are shown for time points at 100 ns and 240 ns where the interaction swaps between D371 and D153. **F**. These dynamics are completely lost in the mutant; in the MD simulation for the D371G mutant, the D367-D153 interaction remains constant throughout the 500 ns window of observations. The small shrinkage in the TCA with the introduction of the mutation is shown in the inset.

Also, the average TCA of the mutant has a narrower distribution than the wild type of closed form which implies other dynamical interactions are involved in the gating mechanism (**Figure 6F, inset**). The entry point has a triangular set of interactions that includes the salt bridge between R367 on the loop joining H7/H8 and D153 on H4 (**Figure S2**). In addition, the nearby D371 on H8 has the potential to form salt bridges with R367 during the dynamical changes that take place in the background fluctuations of the drug free channel as well as during the passage of drugs. The sensitivity conferring mutations at the periplasmic entry observed in all drugs (**Figure 5B**) have the effect of disrupting these interactions between the effluxed molecule and TolC. Introducing the mutation D371G removed a negatively charged residue that competed with carbenicillin for the interaction with R367 and resulted in a more permanent salt bridge with D153. Recalling that the TCA fluctuated less in this region, we find that this rigidity allowed for a larger volume to accommodate the drug molecule. Paradoxically, this increased the residence time of the drug in the initial trapping region because the dynamical hand-over-hand motions of the drug movement were altered. Moreover, this effect impacted the entire efflux pump (**Figure 6C**) and led to the formation of fewer hydrogen bonds along the whole path. We thus conjecture that the D371G mutation caused an allosteric occlusion effect that changed the efflux pattern of the carbenicillin molecule at the extracellular site.

## Discussion

TolC is an outer membrane protein involved in several cellular functions, such as efflux of antibiotics, bile salts, dyes and chemicals, entry of colicin and phages, and haemolysin secretion. Its efflux efficiency is highly dependent on the molecules being pumped out. Here we focus on three β-lactam antibiotics that are effluxed by the TolC at varying rates. The SMD simulations corroborate these findings by quantifying the work done during the efflux and pinpointing interactions that lead to barriers along this path. Moreover, the single residues selected by our experimental screen shed further light on the mode of efflux of each drug molecule. For example, movement of piperacillin is mostly retarded in the β barrel buried in the lipid bilayer near the exit site, while movement of carbenicillin is facilitated both by the hydrophobic plug in the entrance and the hydrophobics lining the first half of the exit pathway (**Figure 4**). In fact, our experimental mutational screen revealed that mutations in the first ∼40 Å region confer increased sensitivity to TolC through several major mechanisms because decreased efflux of antibiotics results in higher effective concentrations of antibiotic molecules inside the cells. These mechanisms include direct interference with dynamical TolC-drug interactions during passage to increase the residence time of the drugs, especially at the entry point. Once an antibiotic enters the channel, the inherent movements of TolC assist in transporting the drug through the majority of the channel. Since TolC collective motions are fine tuned for efflux even in the absence of drugs (**Supplementary Movies S1-S3**), any free molecule in this space has the tendency to be carried towards the extracellular side. The rigidity maintained by the equatorial region is possibly the facilitator of this efficient efflux mechanism, since we found that key mutations in this region significantly alter the efflux. Near the exit, interactions between drug and polar/charged TolC residues form a final barrier before the drug exits the pump. The exit itself is guarded by a hydrophobic patch of residues, which may serve as to gate the entry of pathogens on the extracellular side. Altering this region, such as that by the A269R mutation, apparently disrupts the interactions at the exit; the substantially lowered fitness due to this change might be due to an uncontrolled opening of the otherwise well-protected exit that makes the channel leaky in both directions.

Our mutational scanning experiments and MD simulations demonstrate that TolC is an optimally designed molecular machine for efflux. Our findings indicate that evolutionary selection promotes both local interactions and long-range modifications in the system, which in turn augment efflux efficiency and maximize bacterial survival. All selected mutations manipulate the network of interactions that work by temporary release of key salt bridges to move the drugs along in a hand-over-hand type of motion. More importantly, the effects do not stay local and shifts in interactions at one end are conveyed to the other end via intrinsic motions of this pump. This so-called allosteric occlusion effect between entry and exit regions is again a result of the intricate network of interactions. Our work paves the way to use standardized pulling simulations (SMD or similar) to test the effectiveness of newly designed drugs in efflux pumps. Our findings suggest that drugs designed to manipulate interactions in the entry region of TolC have the best chance of circumventing resistance developed during efflux.

## MATERIALS AND METHODS

### Molecular Dynamics Simulations

The closed state x-ray structure (PDBID: 1EK9)(Koronakis et al. 2000), and the open state cryo-electron microscopy (cryo-EM) structure (PDBID: 5NG5) (Wang et al. 2017) were used as the initial TolC coordinates for MD simulations. The D371G mutant which we hypothesized destroyed key salt bridges on the periplasmic entrance side of TolC was also investigated via MD simulations; the initial coordinates of this mutant were created using the Pymol(Schrödinger LLC 2015). These coordinates were subjected to MD simulations to understand the time dependent behavior of the protein under physiological conditions. Since TolC is a Gram-negative bacterium protein spanning the outer membrane and the periplasmic space, the orientation of TolC was arranged according to the Orientations of Proteins in Membranes (OPM) database (Lomize et al. 2012; Lomize et al. 2022). The overall system was prepared on the CHARMM-GUI platform (Jo et al. 2007; Jo et al. 2008; Best et al. 2012; Lee et al. 2016). The protein was placed in a bilayer that contains 512 POPE lipids and solvated with TIP3 water molecules. Na^+^ and Cl^-^ ions were added to fulfill the isotonic condition of physiological environment (0.15 M).

MD simulations were performed using NAMD and CHARMM36 force field parameters with periodic boundary conditions (Best et al. 2012). The total number of atoms for all our simulations are around 150,000 with box dimensions of 94 x 94 x 173 Å. The particle-mesh Ewald method with a grid spacing of maximal 1 Å per grid in each dimension was used (Essmann et al. 1995). 12 Å cutoff radius with a switching function turned on at 10 Å was applied to calculate van der Waals (vdW) interactions. The RATTLE algorithm (Andersen 1983) was utilized to set the step size of 2 fs during numerical integration with the Verlet algorithm. Temperature regulation was managed with Langevin using a damping coefficient of 5 ps^−1^. Pressure regulation was achieved through a Nose-Hoover Langevin piston, and volume variations were set to remain uniform in all directions. Systems were minimized for 50,000 steps to remove undesired van der Waals contacts. The energy-minimized systems were subjected to 5 ns MD run with constant temperature control (*NVT*). The production parts of the MD simulations of length 500 ns were run in the isothermal-isobaric (*NPT*) ensemble at 310 K and 1 atm. The data from MD simulations were analyzed by the open-source Python package ProDy (Bakan and Bahar 2009; Bakan et al. 2011; Bakan et al. 2014; Zhang et al. 2021) as well as custom Python and tcl scripts.

### Principal Component Analysis (PCA)

Our MD simulations cover the sub-μs time scale dynamics of the protein. Normal mode analysis (NMA) models protein as a harmonic oscillating system and determines the protein dynamics around an energetically stable state of the protein. Low-frequency vibrations, also known as low-energy modes, are associated with collective motions, whereas higher-frequency modes are linked to local deformations (Case 1994; Brooks et al. 1995; Henzler-Wildman, Thai, et al. 2007; Henzler-Wildman, Lei, et al. 2007; Bahar et al. 2010; Skjaerven et al. 2011; Bauer et al. 2019). Principal component analysis (PCA) is an approach for comprehending protein dynamics that involves analyzing an ensemble of conformations obtained from MD simulations (Jolliffe 2002). The mathematical principles underlying PCA and NMA share similarities: both require solving linear equations. In PCA, the variance-covariance matrix (**C**) is diagonalized; **C** approximates the inverse of the Hessian matrix employed in NMA (Bahar et al. 2010). The covariance matrix is constructed using the analysis of multiple snapshots from a MD trajectory, rather than through the calculation of the second derivative near a selected energy minimum. In both instances, the outcomes provide insights into the potential motions a particular structure can undergo, while in a stable conformation. Reducing the dimensions obtained from MD simulations can be advantageous for an unbiased global analysis and to isolate specific configurations. We used principal components to discern the dominant motions of TolC during the MD simulations. PCA was carried out using ProDy (Bakan and Bahar 2009; Bakan et al. 2011; Bakan et al. 2014; Zhang et al. 2021) and the C_α_ atoms were subjected to analysis. In our previous work we have shown that 40 ns chunks of equilibrated trajectories can sufficiently sample the elastic motions around a selected energy minimum of a protein.^52^ We have therefore utilized 240-280 and 460-500 ns fragments of the MD trajectories. Extracellular loops and a part of the equatorial domain which displays large motions obscuring the motions of the overall TolC were excluded in the construction of the **C** matrix.

### Perturbation Response Scanning (PRS)

The PRS method is a powerful tool we previously developed to identify key residues that are functional in the interconversion of one conformer of a protein into another. PRS uses linear response theory to shed light on the possible conformational changes of protein might undego by applying external forces on residues *in silico* (Atilgan and Atilgan 2009). In previous studies (Penkler et al. 2017; Jalalypour et al. 2020), we and others used PRS to study water-soluble proteins. However, TolC is a transmembrane protein with much larger size and requires embedding the TolC homotrimer in a lipid bilayer. Studying TolC dynamics via PRS also provided us the opportunity to evaluate the utility of this powerful method on membrane proteins and solve possible problems arising from this relatively complex environment.

PRS requires experimentally obtained initial and target coordinates of the C_α_ atoms of a protein, **S**_initial_ and **S**_target_, respectively. In our case, these structures were the closed and open forms of TolC. **R**_0_ represents the unperturbed state coordinates (initial), **R**_**1**_ represents the perturbed state coordinates (target) of the C_α_ atoms. The external forces vector (Δ**F)** in random directions are sequentially applied on each residue of the initial structure. Each predicts displacements in the initial structure, Δ**R**_**1**_ by

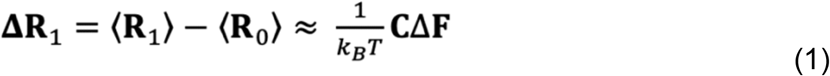

where *k*_B_ is the Boltzmann constant, and *T* is temperature in Kelvin units.

A total of 250 random forces are applied to each C_α_ atom in the protein and results in 250*N* predicted displacement vectors, where *N* is the number of residues. The agreement between experimentally observed and predicted displacements of each residue *i* in response to the applied force is calculated by Pearson correlation coefficient (*C*_*i*_) between predicted and experimental displacements. Thus, each Δ**R**_**1**_ for applied random forces is compared with the experimentally determined difference between **S**_initial_ and **S**_target_ coordinates (Δ**S**), averaged over all affected residues *k*:

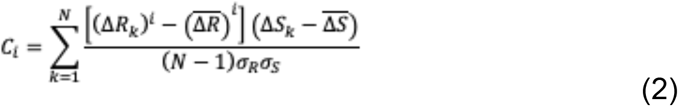

Δ**S**_*k*_ is the experimental displacement between initial MD frame and the target conformation where 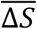 the overbar represents average displacement, and σ_R_ and σ_S_ denote the standard deviations of experimental and predicted structures, respectively. In this equation, the magnitudes of the displacements, *ΔR*_*k*_, are compared with *ΔS*_*k*_. *C*_*i*_ takes on values in the range [0,1] where a correlation greater than 0.7 refers to strong correlation with the experimentally determined conformational changes.

### Steered Molecular Dynamics (SMD)

We performed SMD calculations using the closed-form TolC structure (PDB ID: 1EK9) as the initial point (Koronakis et al. 2000). In separate SMD runs, carbenicillin, piperacillin, or oxacillin was positioned at the periplasmic tip of the protein by docking, CB-Dock2 tool (Liu et al. 2022). The charges of carbenicillin, piperacillin, and oxacillin in water are -2, -1, and -1, respectively. While these charges might shift depending on the position of the drugs in the protein environment, we assumed the pKa of all positions and the charges remained constant in our simulations. The simulation boxes for the membrane, protein, and three different antibiotic molecules were optimized using the CHARMM-GUI web server as described in the “MD simulations” section. Force fields for the antibiotic molecules used in our study were generated by CHARMM general force field module of CHARMM-GUI (Jo et al. 2008; Brooks et al. 2009; Kim et al. 2017). The systems obtained with the three different antibiotics were minimized for 50,000 steps and equilibrated for 0.5 ns. A 50 kcal mol^-1^ Å^-2^ harmonic spring constant was applied to a carbon atom on the hexane structure of antibiotic molecules (depicted as purple spheres in **Figure 1C**) for forcing the drug molecule pass through the TolC channel, a value that satisfies the stiff spring approximation (Nategholeslam et al. 2017). Antibiotics were pulled along the z-axis until they reached the extracellular loops of the protein at a constant velocity of 5 Å ns^-1^. The molecules traveled approximately 140 Å; each SMD simulation thus lasts nearly 30 ns. No restraints were applied to the substrates in the *x* and *y* directions. The pulling simulations of each antibiotic molecule were repeated 45 times to increase the signal-to-noise ratio. The average work done to reach the extracellular side was then calculated for each antibiotic molecule. We note that the general trends and the energy levels in the work curves did not change when the pulling atom was modified to the nitrogen atom in the middle of the CBC molecule from the carbon atom on the hexane ring. Also, modification of the pulling velocity to 0.5 Å ns^-1^ from 5 Å ns^-1^ did not change the profiles in the first 30 Å of the pump.

### Bacterial Strains and Growth Conditions

The BW25113 *E. coli* strain (CGSC No.: 7636) and the *ΔtolC732::kan E. coli* strain (CGSC No.: 11430) were obtained from the Coli Genetic Stock Center. Using the method from reference (Datsenko and Wanner 2000), we removed the kanamycin resistance marker in the *ΔtolC732::kan E. coli* strain. In the article, we refer to the *E. coli* strain *tolC* gene deletion as *ΔtolC* strain. We performed Illumina whole-genome sequencing to inspect potential mutations in these strains, and, apart from the *tolC* deletion, no other mutations were detected (Tamer et al. 2021). Bacterial cells were grown at 37°C in a M9 minimal medium supplemented with 0.4% glucose and 0.2% amicase.

The *tolC* gene from BW25113 (wild type, WT) strain was PCR amplified using 5’-TACCCGGCAGATCTTTGTCGATCCTA-3’ (forward), and 5’-GTGAGCTGAAGGTACGCTGTATCTCA-3’ (reverse) primers. PCR amplified *tolc* gene was then cloned into the pSF-Oxb14 plasmid which was obtained from Oxford Genetics (OGS557, Sigma) using the NEBuilder HiFi DNA Assembly kit (New England Biolabs); by strictly following the protocols recommended by the manufacturers. The resulting plasmid (pSF-Oxb14-t*olC*) contains kanamycin resistance cassette and an Oxb14 constitutively open promoter region. The introduced E. coli *tolc* gene sequence was confirmed by Sanger sequencing. Whole gene saturation mutagenesis library (SML) for each codon in the *tolC* gene used in our experiments was previously prepared in the Toprak lab as described before (Tamer et al. 2021). The *tolC mutant* library were cloned into the pSF-Oxb14 plasmid were transformed into *ΔtolC* strain for downstream selection experiments. We refer to this library as TolC-SML. We refer the *ΔtolC* strain supplemented *with the* pSF-Oxb14-t*olC plasmid as ΔtolC+*p*tolC*. All selection experiments were conducted with minimal M9 media containing 50 μg/ml of kanamycin to prevent loss of the pSF-Oxb14-t*olC plasmid*.

### Determination of minimum inhibitory concentrations (MIC). Bacterial cells were

overnight grown and optical densities (OD600) of cultures were measured in a 1 cm pathlength cuvette. OD600 was then adjusted to 0.001 for which corresponds ∼5×10^5^ colony forming unit (CFU) per milliliters. Bacterial cells were then grown in 96 well plates (∼200 μl per well) in which drug gradients were created using serial dilutions (**Figure 2A**). Highest starting antibiotic concentrations on these plates were 100 μg/ml for carbenicillin, 20 μg/ml for piperacillin, and 1875 μg/ml for oxacillin. The dilution factor for carbenicillin and piperacillin was 1/2 while it was 1/5 for oxacillin. These 96-wells were incubated for 18 hours in a humidity-controlled shaker operated at 37°C and 400 RPM. After the overnight incubation, cell densities were measured using a plate reader (VictorX, PerkinElmer). The experiments were performed in triplicates for each antibiotic molecule. For each strain and antibiotic compounds, the MIC value was defined as the lowest antibiotic concentration at which the final OD600 was below ∼0.04 after background correction.

### Selection Assay

We used three antibiotics: carbenicillin, piperacillin, and oxacillin as selection agents. We used the saturation mutagenesis library (SML) for *tolC* that we previously reported in a publication (Tamer et al. 2021). In this library, all residues except start codon (471 residues in the mature TolC protein, and the 21 residue-long signal sequence) were mutated to all other possible residues, and a pool of mutant library was generated (**Figure 7**). We measured the fitness effects of mutations under selection using a liquid-based sequencing assay. We grew *the* mutant library in M9 minimal medium supplemented with 0.4% glucose and 0.2% amicase overnight, diluted to a final optical density of 0.001, and then exposed these cultures to one of the selection factors: carbenicillin, piperacillin, and oxacillin for 3-hour at 37°C, 400 rotation per minute (RPM). In parallel, we grew cultures without any selection (untreated: UT) for separating drug-induced fitness effects from other potential fitness effects that are not related with the bactericidal effects of antibiotics. The concentrations for these antibiotics in this selection assay were 20 μg/ml and 10 μg/ml for carbenicillin, 10 μg/ml for piperacillin, and 460 μg/ml for oxacillin. These concentrations were close to the MIC of the related compound (**Figure 2B-D**). At the end of the selection period, these cultures were spun down at 5000xg for 5 minutes. The pellets obtained from spinning were resuspended in fresh M9 minimal medium and incubated at 37°C, 400 RPM for 6 hours to increase cell density and ensure harvesting of enough *tolC* carrying plasmid for sequencing. We recorded cell densities during this recovery period as depicted in **Figure S1**. Cells were harvested by spinning cultures at 5000xg for 5 minutes. Pellets were collected for plasmid purification. *tolC* gene variants on these plasmids were amplified with polymerase chain reaction (PCR) using 5’-TACCCGGCAGATCTTTGTCGATCCTA-3’ (forward), and 5’-GTGAGCTGAAGGTACGCTGTATCTCA-3’ (reverse) primers.

**Figure 7.**
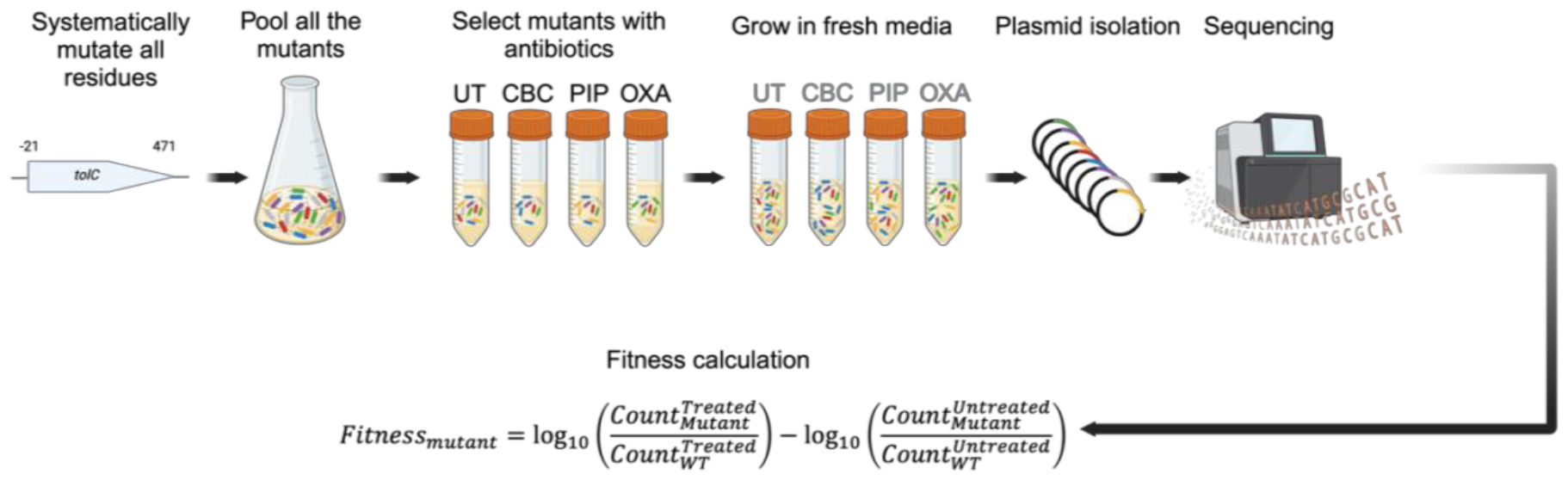
Selection assay protocol. We conducted deep mutational scanning on TolC, systematically mutating all 493 residues, which includes the 21-amino acid long signal sequence preceding the start codon (excluding it). Residues were mutated to all the 19 other amino acids, along with the stop codon. Obtained mutants were pooled together and grown overnight. Overnight grown mutant pool and *ΔtolC* strains were split into cultures: untreated (UT), CBC, PIP, OXA; the experiment was replicated once. Cultures other than the untreated were treated with the relevant selection factors. Cultures were exposed to drugs for 3 hours for selection. This was followed by 6 hours long incubation in fresh growth media without any selection agent for the recovery of cells (gray abbreviations on the tubes represent previously applied antibiotics). We isolated *tolC* mutant carrying plasmids and PCR amplified the tolC gene for amplicon sequencing. Mutation frequencies were calculated by deep sequencing and fitness values of each mutation were calculated using the formula in the figure.

These products were sequenced on Illumina NovaSeq 6000, producing 151 bp reads. Sequence reads were compared with the wild type *tolC* sequence and mutations were listed. Sequence reads that had mutations in more than one residue were excluded from the analysis. Synonymous mutations yielding the same amino acid replacement were grouped together and used as a reference for relative fitness calculations. The frequency of each mutation was calculated by dividing the number of counts for that mutation with the number of all reads, including alleles with multiple mutations.^21^ We then used the formula displayed in **Figure 7** to calculate relative fitness effects of each mutation.

## Acknowledgements

The numerical calculations reported in this paper were partially performed at TUBITAK ULAKBIM, High Performance and Grid Computing Center (TRUBA resources). We thank TUBITAK project no. 122F149 for partial support. IK was partially supported by TUBITAK 2214-A visiting research fellowship (Application Number: 1059B142100453). E.T. is supported by UTSW Endowed Scholars Program, Human Frontiers Science Program Research grant RGP0042/2013, NIH grant R01GM125748, DOD PR172118, and Welch Foundation I-2082-20210327.

## Data and Software Availability

Raw amplicon sequencing data for the mutagenesis experiment are deposited under the NCBI BioProject with accession number PRJNA1078904. Original code generated for producing the results for this study is deposited at GitHub at the following repository: https://github.com/midstlab/Kantarcioglu_2024. Any additional data provided in this paper will be provided by the lead contact upon request.

