## Supporting Information for "Elucidating TolC Protein Dynamics: Structural Shifts Facilitate Efflux Mediated β-lactam Resistance"

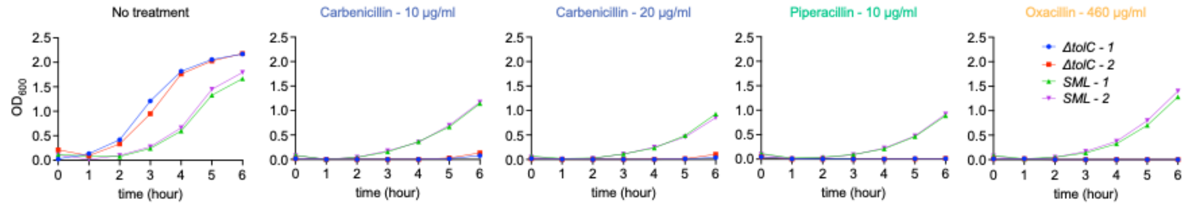


**Figure S1 | Growth curves after the selection agent incubation.** After the 3-hour selection agent incubation, surviving cells from *ΔtolC*, *SML*, and their replicate cultures were transferred to fresh M9 minimal media. The growth of these cells was monitored for 6 hours. Cultures for no treatment case were grown properly. *SML* cultures containing plasmids in their cells started to grow properly after 3 hours of incubation in fresh media. In contrast, *ΔtolC* cultures did not show growth within the represented period, suggesting the importance of the *tolC* gene for survival in the presence of these antibiotics. Blue and red colors represent *ΔtolC* cultures and green and magenta colors represent *SML* cultures.

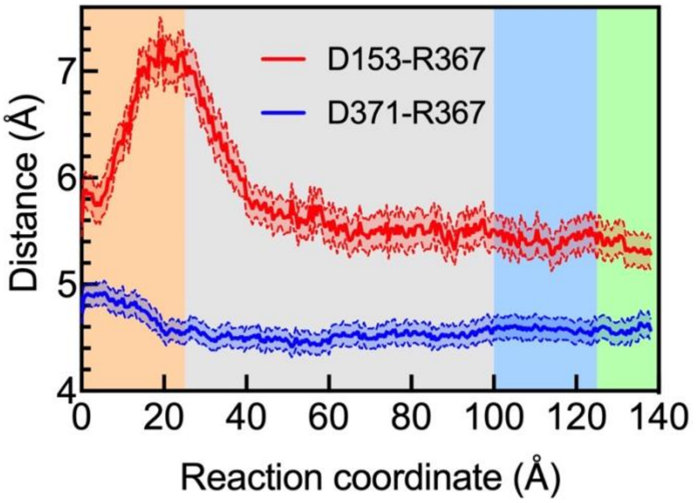


**Figure S2 | Salt bridge distance between R367 and D371 during carbenicillin passage in the wild type.** The average donor-acceptor distance for R367 with D371 and D153 residues show that the salt bridge is disrupted until carbenicillin moves past the first 30 Å and is then reestablished. Averages of 45 SMD simulations and the three chains in TolC are shown with SEM.

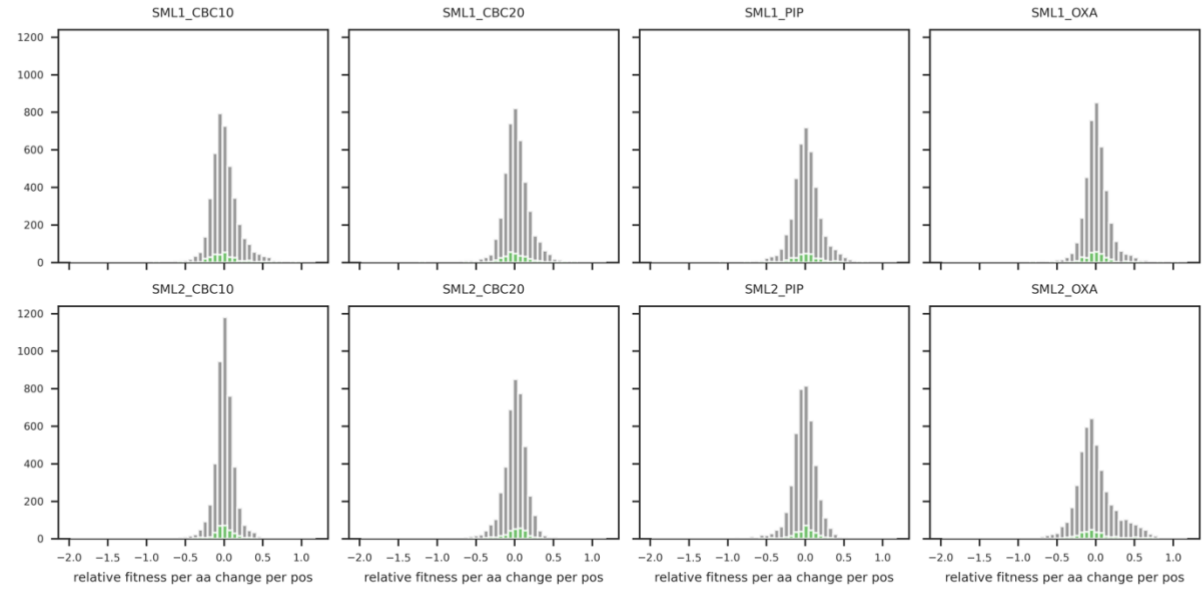


**Figure S3 | Relative fitness histograms.** Grey histograms show the distribution of relative fitness measurements of all amino acid changes detected in the corresponding sample and replica. Green overlayed histograms show the distribution of relative fitness measurements of synonymous mutations. Relative fitness is defined as the log10 value of fraction of a mutant allele in the drug treated sample divided by the untreated.

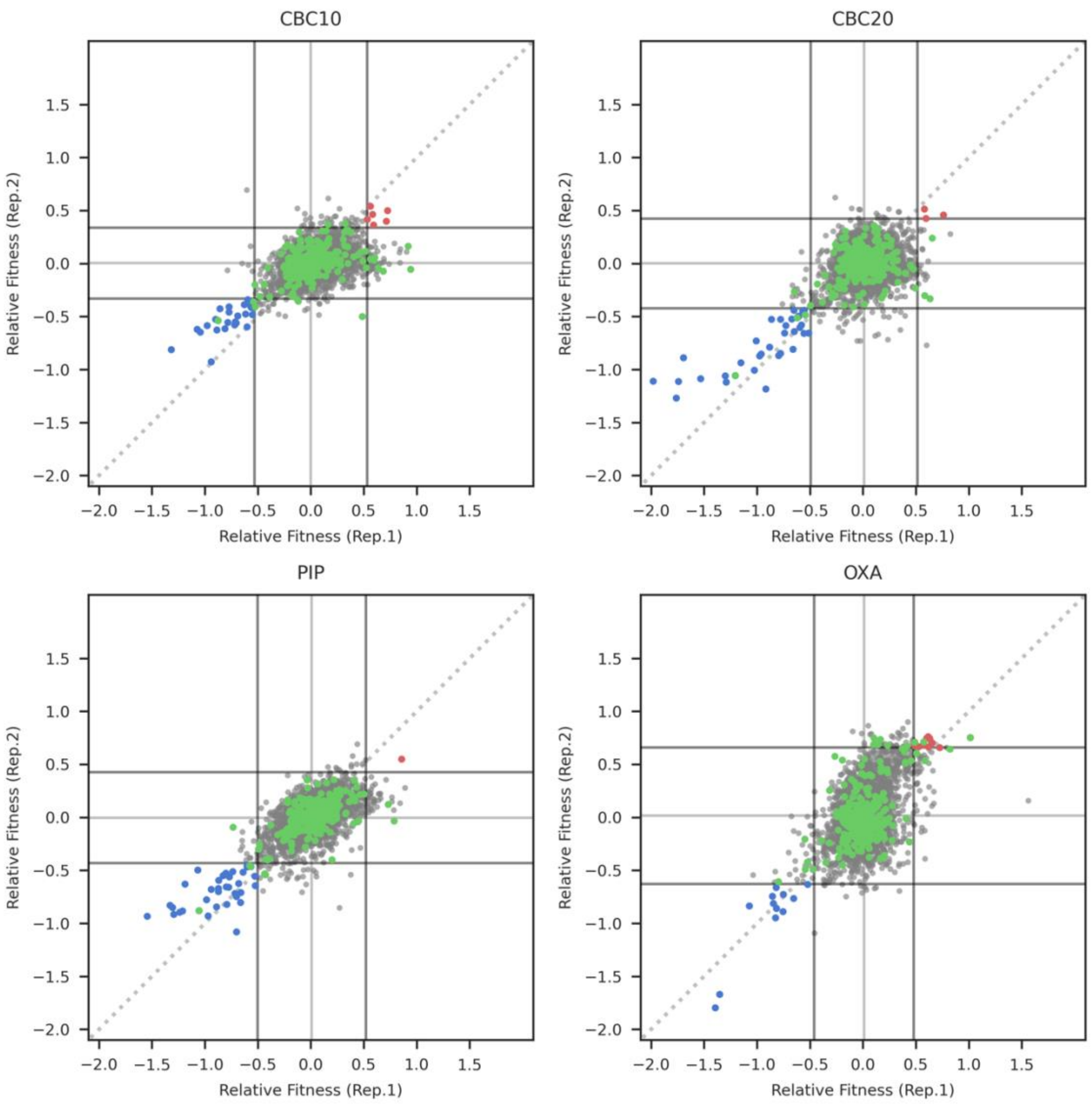


**Figure S4 | Significance tests at 2.5σ of both samples (99.4 % confidence interval).** Scatter plots show the relative fitness measurements of the same amino acid changes in two biological replicates. Grey dots represent all mutant alleles and green dots represent synonymous mutant alleles. A significance threshold of 2.5 standard deviations of synonymous mutation relative fitness distribution in both replicates is used to determine significantly resistance and sensitivity conferring mutations, indicated in red and blue, respectively.

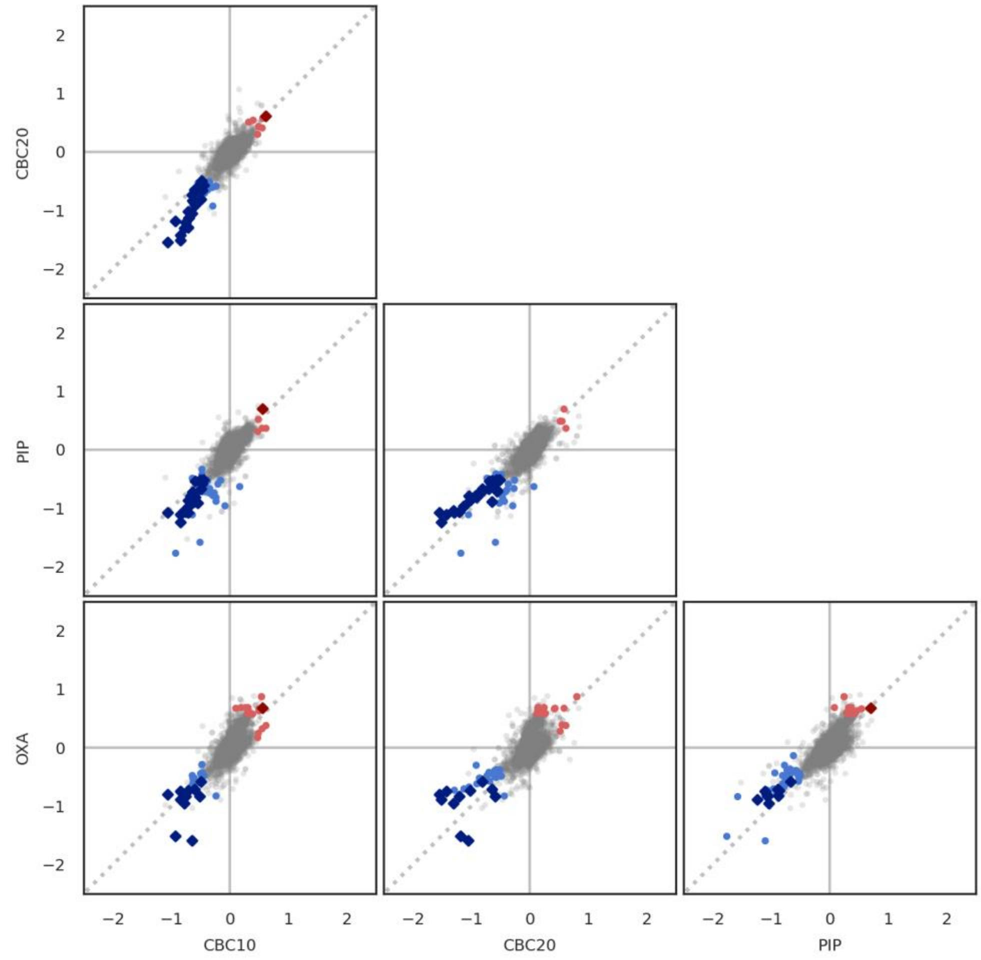


**Figure S5 | Drug pair significance tests.** Scatter plots show the relative fitness measurements of the same amino acid changes across different drug selections. Mean relative fitness of two biological replicates used to represent each mutant allele. Grey dots represent all mutations, bright red dots represent mutants significantly resistance conferring in at least one of the drugs, bright blue dots mutants significantly sensitivity conferring in at least one of the drugs, and dark diamond markers show mutants that are significantly resistance/sensitivity conferring in both drugs.

### Table S1 | Minimum inhibitory concentration data used to construct figure 2.

| CBC 100 µg/ml | *WT* | | | *ΔtolC* | | | *WT ΔTolC+ oxb14 tolC* | | | *SML* | | |
|---|---|---|---|---|---|---|---|---|---|---|---|---|
| 100 | 0 | 0.002 | 0.002 | 0 | 0.001 | 0.002 | 0.001 | 0.002 | 0.002 | 0.001 | 0.001 | 0.002 |
| 50 | 0.001 | 0.002 | 0.002 | 0.001 | 0.001 | 0.002 | 0.001 | 0.002 | 0.002 | 0.001 | 0.001 | 0.002 |
| 25 | 0.001 | 0.002 | 0.002 | 0.001 | 0.002 | 0.001 | 0.001 | 0.002 | 0.002 | 0 | 0.001 | 0.002 |
| 12.5 | 0.002 | 0.02 | 0.003 | 0.001 | 0.002 | 0.002 | 0.002 | 0.002 | 0.003 | 0.001 | 0.002 | 0.002 |
| 6.25 | 0.002 | 0.003 | 0.003 | 0.001 | 0.002 | 0.002 | 0.002 | 0.002 | 0.002 | 0.001 | 0.002 | 0.072 |
| 3.125 | 0.356 | 0.359 | 0.272 | 0.001 | 0.002 | 0.001 | 0.002 | 0.002 | 0.002 | 0.001 | 0.042 | 0.117 |
| 1.5625 | 0.326 | 0.336 | 0.334 | 0.001 | 0.002 | 0.002 | 0.277 | 0.261 | 0.263 | 0.078 | 0.06 | 0.109 |
| 0.78125 | 0.398 | 0.573 | 0.489 | 0.283 | 0.337 | 0.315 | 0.154 | 0.168 | 0.135 | 0.145 | 0.156 | 0.153 |
| 0.390625 | 0.599 | 0.573 | 0.567 | 0.454 | 0.44 | 0.422 | 0.237 | 0.225 | 0.202 | 0.214 | 0.252 | 0.263 |
| 0.1953125 | 0.589 | 0.588 | 0.581 | 0.448 | 0.436 | 0.403 | 0.232 | 0.222 | 0.203 | 0.257 | 0.266 | 0.27 |
| 0.09765625 | 0.587 | 0.544 | 0.544 | 0.446 | 0.41 | 0.393 | 0.245 | 0.225 | 0.213 | 0.269 | 0.256 | 0.278 |
| 0 | 0.562 | 0.567 | 0.563 | 0.417 | 0.396 | 0.402 | 0.258 | 0.227 | 0.214 | 0.282 | 0.284 | 0.284 |

| PIP (20µg/ml) | *WT* | | | *ΔtolC* | | | *WT ΔTolC+ oxb14 tolC* | | | *SML* | | |
|---|---|---|---|---|---|---|---|---|---|---|---|---|
| 20 | 0.005 | 0.004 | 0.004 | 0 | 0.001 | 0.001 | 0.001 | 0.001 | 0.002 | 0.002 | 0.003 | 0.003 |
| 10 | 0.004 | 0.004 | 0.005 | 0.002 | 0.002 | 0.002 | 0.001 | 0.002 | 0.002 | 0.003 | 0.003 | 0.004 |
| 5 | 0.003 | 0.005 | 0.005 | 0.001 | 0.001 | 0.002 | 0.001 | 0.002 | 0.002 | 0.004 | 0.003 | 0.003 |
| 2.5 | 0.003 | 0.005 | 0.005 | 0.001 | 0.002 | 0.002 | 0.001 | 0.002 | 0.003 | 0.003 | 0.003 | 0.003 |
| 1.25 | 0.35 | 0.316 | 0.374 | 0.002 | 0.002 | 0.002 | 0.049 | 0.06 | 0.027 | 0.146 | 0.005 | 0.188 |
| 0.625 | 0.372 | 0.399 | 0.408 | 0.001 | 0.002 | 0.001 | 0.174 | 0.179 | 0.157 | 0.198 | 0.186 | 0.226 |
| 0.3125 | 0.597 | 0.605 | 0.621 | 0.002 | 0.002 | 0.002 | 0.264 | 0.309 | 0.286 | 0.278 | 0.267 | 0.284 |
| 0.15625 | 0.614 | 0.613 | 0.603 | 0.002 | 0.002 | 0.002 | 0.268 | 0.322 | 0.303 | 0.305 | 0.303 | 0.309 |
| 0.078125 | 0.601 | 0.587 | 0.594 | 0.002 | 0.002 | 0.002 | 0.278 | 0.324 | 0.317 | 0.307 | 0.31 | 0.313 |
| 0.0390625 | 0.605 | 0.609 | 0.612 | 0.01 | 0.046 | 0.349 | 0.299 | 0.317 | 0.331 | 0.305 | 0.316 | 0.357 |
| 0.01953125 | 0.598 | 0.562 | 0.59 | 0.375 | 0.383 | 0.392 | 0.298 | 0.285 | 0.327 | 0.342 | 0.33 | 0.37 |
| 0 | 0.55 | 0.551 | 0.543 | 0.4 | 0.38 | 0.411 | 0.334 | 0.326 | 0.344 | 0.339 | 0.358 | 0.359 |

| OXA (1875 µg/ml) | *WT* | | | *ΔtolC* | | | *WT ΔTolC+ oxb14 tolC* | | | *SML* | | |
|---|---|---|---|---|---|---|---|---|---|---|---|---|
| 1875 | 0.001 | 0.001 | 0.064 | 0.0001 | 0.0001 | 0.0001 | 0.0001 | 0.0001 | 0.0001 | 0.02 | 0 | 0 |
| 375 | 0.225 | 0.294 | 0.279 | 0.0001 | 0.0001 | 0.0001 | 0.004 | 0.003 | 0.01 | 0.001 | 0.06 | 0.001 |
| 75 | 0.577 | 0.558 | 0.567 | 0.001 | 0 | 0 | 0.31 | 0.323 | 0.306 | 0.27 | 0.26 | 0.268 |
| 15 | 0.576 | 0.588 | 0.555 | 0.002 | 0 | 0 | 0.295 | 0.313 | 0.267 | 0.286 | 0.278 | 0.305 |
| 3 | 0.589 | 0.597 | 0.572 | 0 | 0.003 | 0 | 0.285 | 0.256 | 0.246 | 0.286 | 0.273 | 0.304 |
| 0.6 | 0.579 | 0.57 | 0.58 | 0 | 0 | 0.001 | 0.247 | 0.261 | 0.224 | 0.275 | 0.288 | 0.308 |
| 0.12 | 0.598 | 0.595 | 0.593 | 0.388 | 0.393 | 0.395 | 0.255 | 0.245 | 0.217 | 0.282 | 0.285 | 0.317 |
| 0.024 | 0.585 | 0.6 | 0.573 | 0.392 | 0.395 | 0.411 | 0.223 | 0.25 | 0.201 | 0.28 | 0.287 | 0.318 |
| 0.0048 | 0.576 | 0.587 | 0.572 | 0.388 | 0.388 | 0.4 | 0.213 | 0.245 | 0.209 | 0.288 | 0.298 | 0.324 |
| 0.00096 | 0.592 | 0.588 | 0.589 | 0.396 | 0.405 | 0.427 | 0.201 | 0.202 | 0.202 | 0.278 | 0.285 | 0.31 |
| 0.000192 | 0.559 | 0.574 | 0.562 | 0.402 | 0.401 | 0.422 | 0.203 | 0.201 | 0.21 | 0.283 | 0.292 | 0.34 |
| 0 | 0.554 | 0.525 | 0.562 | 0.403 | 0.406 | 0.425 | 0.201 | 0.207 | 0.21 | 0.268 | 0.293 | 0.289 |

### Supplementary Videos:

The coloring scheme for protein maintained throughout the manuscript is also valid for Supplementary Videos.

Residues that formed two or more hydrogen bonds for each antibiotic molecule are highlighted as dynamic bonds. Positively charged residues are red, polar residues are violet, and hydrophobic residues are cyan-colored. In addition to these hydrogen bonding residues, R367 is highlighted as red and D371 is highlighted as blue.

**Video S1 |** PC1 obtained from the 460-500 ns portion of the closed trajectory

**Video S2 |** PC2 obtained from the 460-500 ns portion of the closed trajectory

**Video S3 |** PC3 obtained from the 460-500 ns portion of the closed trajectory

**Video S4 |** PC1 obtained from the 460-500 ns portion of the open form trajectory

**Video S5 |** PC2 obtained from the 460-500 ns portion of the open form trajectory

**Video S6 |** PC3 obtained from the 460-500 ns portion of the open form trajectory

**Video S7 |** Sample SMD trajectory displaying the interactions of carbenicillin with the high contact residues in TolC

**Video S8 |** Sample SMD trajectory displaying the interactions of piperacillin with the high contact residues in TolC

**Video S9 |** Sample SMD trajectory displaying the interactions of oxacillin with the high contact residues in TolC
